# Seventy-five years of insight – the impact of the Ukulinga very long-term grassland experiments

**DOI:** 10.1101/2025.05.30.656967

**Authors:** Craig D Morris, Kevin P Kirkman

**Affiliations:** Agricultural Research Council – Animal Production Institute (ARC-API), c/o University of KwaZulu-Natal, Private Bag X01, Scottsville 3209, South Africa; School of Life Sciences, University of KwaZulu-Natal, Private Bag X01, Scottsville 3209, South Africa

**Keywords:** disturbance, fire, fertilisers, mowing, nutrients

## Abstract

Two of the world’s oldest grassland experiments began in 1950–51 in Pietermaritzburg, South Africa. The Burning and Mowing Trial (BMT) tests how summer mowing and dormant-season burning or mowing regimes affect mesic grassland. The Veld Fertiliser Trial (VFT) examines how nitrogen, phosphorus fertilisers and lime influence grassland productivity, composition, and diversity. Seventy-five years later, their impact was assessed through research output and value for education and networking. All published papers and their citations were identified, with key themes, leading authors, and citation patterns of the top ten papers from each trial analysed. A total of 25 and 24 peer-reviewed papers, cited 1,652 times, have been published from the BMT and VFT, respectively, with publication rates rising since 2000. Studies addressed treatment effects on soil, plants, invertebrates, microbes, regional comparisons, and remote sensing. South Africans led authorship, with strong USA participation, and the top-10 papers reached a wide multinational audience. The Ukulinga LTEs demonstrated the vital role of regular burning or mowing, revealed how fertilisers can negatively impact grassland, and highlighted the importance of treatment interactions in shaping ecosystem dynamics. As key sites for research collaboration and education, they should be maintained long-term to advance understanding and address new questions.

## INTRODUCTION

Around 250 years ago, humans unknowingly initiated an unprecedented global experiment by rapidly unleashing fossil fuels that had taken millions of years to accumulate (Weart 2010; Höök and Tang 2013). The burning of coal, oil, and gas during the Industrial Age has emitted vast amounts of CO₂, CH₄, N₂O, and other greenhouse gases, triggering warming and complex climate effects with widespread consequences for ecosystems, sea levels, and human livelihoods (Ledley et al. 1999; Hansen and Stone 2016; IPCC 2022). The existential threat of global climate change—now more urgently called the climate crisis—has driven extensive research to understand its processes, causes, and mitigation strategies (Haunschild et al. 2016). Long-term studies like the Free-Air CO_2_ Enrichment (FACE) experiments (Norby and Zak 2011) play a key role in climate research, simulating elevated CO₂ levels (Casella and Soussana 1997) and warming (Shi et al. 2015) to observe how ecosystems respond and revealing the interactive nature of these responses (De Boeck et al. 2015). Unlike the unplanned global climate perturbation, where ‘treatments’ are unreplicated, shifting, and intensifying (Wu et al. 2015; Stoddard et al. 2021), long-term studies like FACE apply controlled, consistent interventions to uncover ecological cause and effect.

The value of long-term experiments (LTEs), such as those used in climate change and agroecological studies, lies primarily in their fixity and longevity. For instance, J.B. Lawes’s decision in 1843 to apply fixed treatments and grow single, non-rotating crops on each plot made the classical LTEs at Rothamsted, UK—still running today—uniquely valuable for understanding fertiliser impacts on individual crops (Johnston 1994). A well-designed LTE applies consistent treatments to test specific hypotheses and build models, with sufficient replication to estimate main and interaction effects precisely (Scheiner and Gurevitch 2001; Wells et al. 2025) and spatial blocking to minimise confounding by environmental variation, especially soils (Barnett 1994; Casler 2018). Long-term treatment maintenance is essential for detecting directional trends, cycles, and successional changes beyond short-term fluctuations caused by rare events (Jentsch et al. 2007; Riginos et al. 2024). Such prolonged experiments can reveal how treatments gradually shape ecosystems and distinguish temporary responses from lasting shifts (Silvertown et al. 2010; Lindenmayer al. 2012; Knapp et al. 2012; Blanc and Thrall 2024). Beyond tracking treatment impacts, LTEs can serve as outdoor laboratories for detecting unforeseen top-down influences (Magnuson 1990; Rasmussen et al. 1998) like climate change (Donmez et al. 2023), nitrogen deposition (Stevens et al. 2022), and unanticipated anthropogenic impacts such as nuclear radioactive fallout (Jenkinson et al. 2008). Baseline samples and ongoing data collection are indispensable for assessing these external factors and their interactions with experimental treatments (Johnston and Poulton 2018). Moreover, LTEs established in different locations enable comparisons across similar or contrasting vegetation types and environments, facilitating collaborative research networks (Vanderbilt and Gaiser 2017) that can uncover common, generalisable, and context-dependent effects, and reveal deeper ecological principles (Knapp et al. 2004; Borer et al. 2017).

In 1950, two long-term experiments were initiated in mesic grassland (veld) dominated by subtropical (C_4_) grasses on the plateau of the Ukulinga Research Farm in Pietermaritzburg, by Professor JD Scott of the newly established Department of Pasture and Soil Science at the University of Natal (now the University of KwaZulu-Natal). These two LTEs are: (1) the Veld Burning and Mowing Trial (BMT), and (2) the Veld Fertilisation Trial (VFT) (See Figure 1 for location). For international conformity and communication, they have since been renamed the Ukulinga Grassland Fire Experiment (UGFE) and the Ukulinga Grassland Nutrient Experiment (UGNE). However, they are referred to here as the BMT and VFT for brevity. Originally designed to address agricultural concerns, the BMT aimed to test the effects of mowing and burning on hay yield and quality, while the VFT focused on assessing the response of grassland herbage yield and quality to various fertiliser treatments. Both experiments have continued uninterrupted for 75 years and are now among the longest-running ecological experiments in the world (Morris and Tainton 2002; Pooley 2018). Their duration qualifies them as “very long-term studies” (20– 50 years) (Viblanc and Muller-Landau 2025).

**Figure 1:**
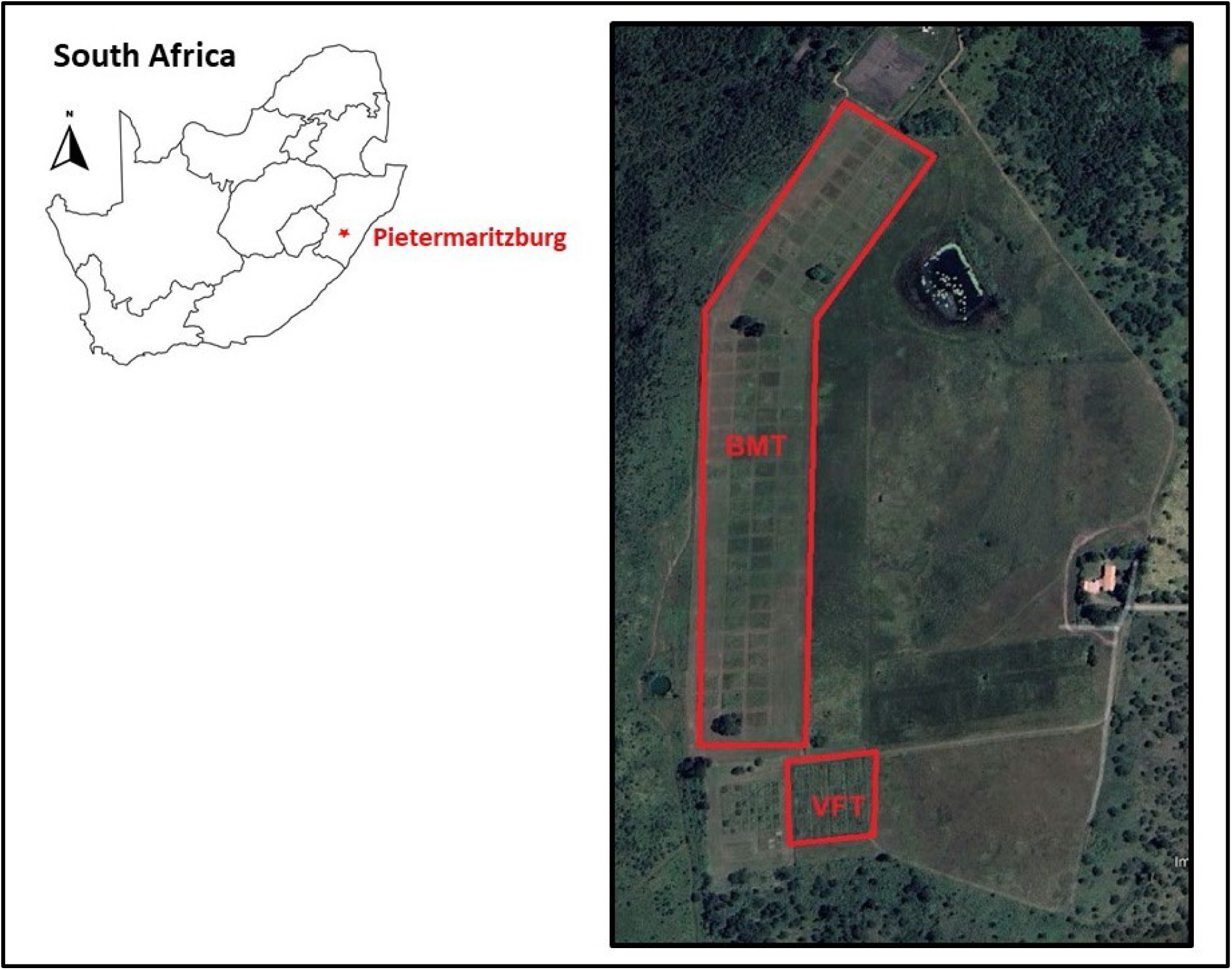
The location of the Burning and Mowing Trial (BMT) and Veld Fertiliser Trial (VFT) on the plateau of the Ukulinga Research Farm, University of KwaZulu-Natal, Pietermaritzburg, South Africa. Coordinates: 29°40’11.18“ S, 30°24’10.83” E

The specific aims of the two Ukulinga LTEs, as specified in the original project proposals, were: (1) for the BMT – to build on earlier experimental work on fire in KwaZulu-Natal (sensu Scott 1947; Staples 1930) in the southern Tall Grassveld, to determine the optimum management strategy in terms of when and how often to burn and defoliate (Rodel 1950; Ions 1960); and (2) for the VFT – to increase productivity from natural grasslands and to provide a more complete picture of the delicate balance that exists between the plant community and its environment. This was based on the recognition of nutrient deficiencies in South African grasslands, namely a lack of phosphate, calcium, and nitrogen (Booysen 1954). However, over time, the focus of the Ukulinga LTEs has also broadened from agriculture to ecology, supporting internationally funded research into grassland ecosystem processes.

The following is a brief account of the historical research developments that led to the establishment of the Ukulinga LTEs.

Since the early 1900s, researchers in South Africa have documented growing concerns about grassland degradation and its negative impacts on livestock production and the national economy (Hall 1934). During this period, controversy emerged surrounding grassland burning practices as land managers and scientists debated optimal management approaches (Staples 1926).

The first rigorous experimental work addressing these issues began at Cedara School of Agriculture in 1921, where researchers established trials investigating the effects of burning, grazing, mowing, and mechanical disturbances on veld condition (Staples 1926). The results proved illuminating—dormant season burning promoted the dominance of *Themeda triandra*, a highly valuable grazing species, while plots without fire and grazing, or with only light grazing, or those subjected to heavy grazing without burning saw this desirable species replaced by less palatable grasses and encroaching shrubs (Staples 1930).

Building on this foundation, further research on burning impacts across various veld types was initiated near Estcourt in KwaZulu-Natal in 1937 (Scott 1951). These experiments compared multiple burning regimes, including unburned controls and various seasonal burns (winter, spring, and autumn) at different frequencies (annual, biennial, and triennial). The research also examined mowing as an alternative to burning. Results indicated that spring mowing without fire maintained the healthiest grass cover, while autumn burning proved particularly harmful. Burning after the first spring rains emerged as the least damaging fire treatment.

Following his work at Estcourt, Scott established a similar but more comprehensive experiment at Ukulinga (Rodel 1950). This experiment—the BMT—was laid out in a split-plot, randomised blocks design (three replications) with four whole-plot treatments and eleven split-plot treatments with plot sizes of 13.7 x 18.2 m. Given the agricultural priorities at the time, the treatments aimed at conserving winter feed by mowing for hay at different times during the growing season. The split treatments were various methods (burning or mowing) of removing undesirable thatch from grassland, which would impede haymaking. Whole-plot treatments were: (A) no mowing or burning for comparison with treated plots, (B) one cut in the early season at an approximate grass height of 200 mm, (C) one cut at the end of February, coinciding with agricultural practice for haymaking, and (D) two cuts, one at time B and one at time C. The split treatments comprised: controls (1) with no removal of old growth, (2) annual winter burns in August, (3) annual spring burns after rain, (4), biennial winter burns, (5) biennial spring burns, (6) biennial burn in autumn, (7) triennial winter burn, (8) triennial spring burn, (9) triennial autumn burn, (10) annual winter mow at the same time as 2, and (11) and annual spring mow at the same time as 3. For the full details on the study site, experiment design and treatments, and layout of the BMT, see Fynn et al. (2004).

Concurrent with these fire management studies, researchers raised concerns about the inherently low forage production and quality of South African veld (Hall 1934; Meredith 1943). Early proposals suggested fertilising natural grasslands to enhance livestock production through increased forage quantity and quality. Pioneering work in this area began at Frankenwald Botanical Research Station north of Johannesburg in 1932, where scientists applied various combinations of nitrogen, phosphorus, potassium, and calcium to natural grassland. The results were promising, showing increased grass yields and improved animal production, along with shifts in grass species composition (Meredith 1948).

These encouraging findings led to the establishment of a comprehensive nutrient addition experiment— the VFT—at Ukulinga Research Farm in 1950 (Booysen 1954). This experiment addressed the recognised deficiencies of phosphorus in veld forage and the limiting effects of nitrogen and low soil pH on productivity. The treatments applied in the VFT included phosphorus (P) at two levels (0 and 28 kg P ha^-1^ year^-1^), lime at two levels (0 and 11 t ha^-1^ every 5 years), and nitrogen at 4 levels (0, 70, 141 and 211 kg N ha^-1^ year^-1^) (Booysen 1954). The N was supplied from two sources, namely ammonium sulphate and ammonium nitrate to investigate the acidifying effect of ammonium sulphate. In later years, the ammonium nitrate was replaced with limestone ammonium nitrate, due to prohibitions on the sale of ammonium nitrate as a fertiliser in South Africa. Plot size was 2.7 x 9 m, and treatments were applied alone and in combination in a 4 x 23 factorial design, with three replications (Barnes et al. 1987). For the full details on the study site, experimental design and treatments, and layout of the VFT, see Zama et al. (2023).

Although long-term field experiments are an irreplaceable cornerstone of ecological and agricultural research (Knapp et al. 2012), their high costs and maintenance needs (Lindenmayer 2018)—often challenging to justify for sustained funding (Lindenmayer et al. 2012; Blanc and Thrall 2024)—mean their scientific, educational, and societal value must be carefully considered (Peterson et al. 2012), especially when underutilised or poorly monitored (Körschens 2006). Therefore, we look back over three-quarters of a century of research from the two long-term grassland trials at Ukulinga to assess the scientific value they have yielded through published output: Has the research been prolific? Has it revealed unique insights that advance the understanding and management of grassland? How far have these findings reached broader audiences, and have they offered wider societal value through formal or informal educational outreach? The specific objectives of our evaluation were to quantify: (1) the output of peer-reviewed scientific papers, (2) the common themes or topics addressed, (3) the distribution of publications across authors and journals, and (4) citation patterns of key papers produced on research conducted on the BMT and VFT. In doing so, we aim to assess the overall scientific value of the Ukulinga LTEs and present an evidence-based case for their continuation (or not) in their current or revised form, ensuring they remain as effective as possible moving forward.

## METHODS

A comprehensive list of all peer-reviewed scientific papers arising from research on the BMT and VFT since their inception was compiled using publication records maintained by the Discipline of Grassland Science, School of Life Sciences, at the University of KwaZulu-Natal (UKZN) in Pietermaritzburg. This was supplemented by input from researchers involved in the trials and additional searches on Google Scholar using keyword combinations such as “Ukulinga,” “grassland OR veld,” and “long-term experiment.”

Information was extracted from the bibliographic details and full texts of each paper to record the primary and co-authors, their countries of affiliation, publication date, journal name, and other relevant details, such as the apparent main topic or theme addressed by each study.

Citation statistics for each paper were sourced from Google Scholar, which captures a broader range of citations, including theses, grey literature, books, conferences, preprints, unpublished reports, as well as the scientific journals indexed by other large scholarly databases, to assess the widest possible reach of the publication (Martín-Martín et al. 2018). Citations of the ten most cited papers from the BMT and VFT in the scientific literature were retrieved from the Scopus bibliographic database, which indexes a wide range of journals (Joshi 2016), along with information on the citing papers’ lead authors, their country affiliations, and the journals in which the Ukulinga LTE papers were cited.

The extracted bibliographic details were summarised using descriptive statistics such as means, medians, and frequencies to compare the scientific output of the two LTEs. Citation patterns were aggregated using the H-index, which represents the highest number of papers (h) from each experiment with at least (h) citations, and the i10-index, which counts the number of papers with at least 10 citations (Noruzi 2016). The Kolmogorov-Smirnov test was used to compare the frequency distribution of citation rates between the two experiments, conducted using Genstat 23 (VSN International 2022).

## RESULTS

### Publication output

A total of 25 and 24 peer-reviewed papers from the BMT and VFT, respectively, have been published in scientific journals since the start of the LTEs (Supplementary Tables S1 & S2). Research output from both experiments was minimal before the turn of the century but increased sharply in the 2000s (Figure 2), with the VFTs’ publication rate initially lagging behind the BMT’s before surpassing it in the 2020s.

**Figure 2:**
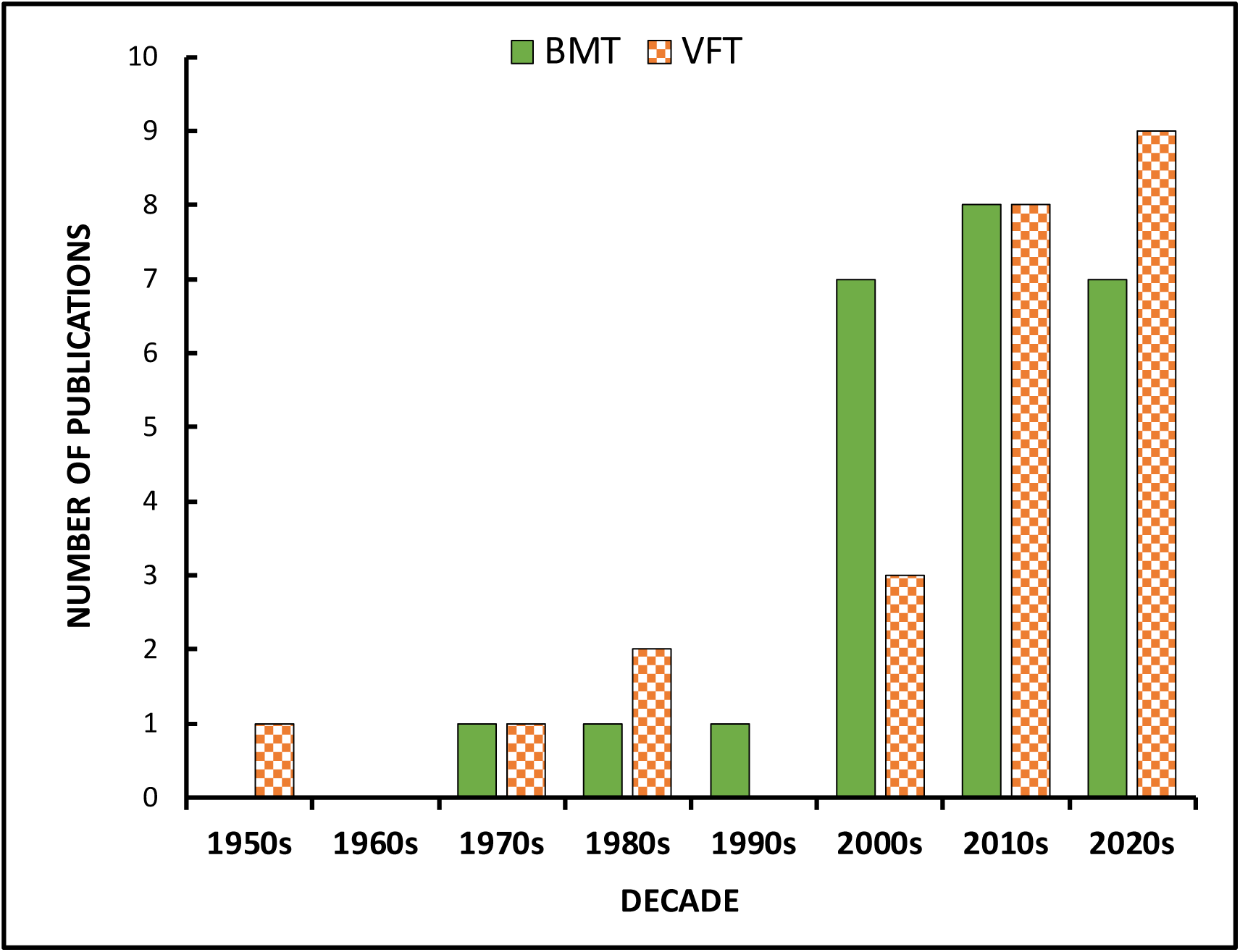
Count of peer-reviewed papers produced in each decade from the Burning and Mowing Trial (BMT) and Veld Fertiliser Trial (VFT) at Ukulinga, South Africa.

The first VFT results were reported by Scott and Booysen (1956) in the South African Journal of Science six years after the experiment began. Fertilisers, especially ammonium nitrate and superphosphate, significantly boosted herbage protein yields, while lime had no effect and nitrogen caused only minor plant composition shifts. By the mid-1980s, le Roux and Mentis (1986) and Barnes et al. (1987) found that nitrogen fertilisers had significantly altered grass species composition and rangeland condition, with phosphate and lime having lesser effects but enhancing the impact of N.

Key interactions between disturbances also emerged in the first BMT results published by Tainton et al. (1978), nearly 30 years after the experiment began. They reported that annual, biennial, or triennial burning reduced herbage yield in the following season but had no lasting effect on productivity. However, veld response varied with summer management, as hay mowing influenced post-burn outcomes. Two focused studies followed, examining how treatment-induced changes in vegetation structure and composition affected the sward’s microclimate (Savage and Vermeulen 1983) and grasshopper communities (Chambers and Samways 1998).

Studies on treatment effects in both LTEs continued into the 2000s, becoming more intensive and multidisciplinary, with an increased focus on plant species diversity (e.g., Fynn et al. 2004; Zama et al. 2023) and a broader consideration of soil (e.g., Fynn et al. 2003; Schleuss et al. 2019) and other biota, including ants (Khoza et al. 2023) and microbes (e.g., Zeglin et al. 2007; Vermeire et al. 2021). The diverse vegetation of the two experimental sites also offered a unique opportunity to test specialised remote sensing methods (e.g., Sibanda et al. 2017a; Naicker et al. 2024). Research expanded to explore plant and soil ecology, using cross-experimental comparisons between the VFT, BMT (e.g., Ward et al. 2020), and other subtropical and temperate grasslands to identify unique and shared ecological responses from findings in similar long-term grassland experiments outside Africa (e.g., Swemmer et al. 2007; Buis et al. 2009; Gold et al. 2023).

### Publication themes

Tables 1 and 2 group LTE publications by their primary topic focus, though most studies are multifaceted and address multiple topics beyond their main focus.

**Table 1:** Papers produced from the BMT grouped by general topic.

| TOPIC | STUDIES | NO. |
| --- | --- | --- |
| **Effects on soil and ecosystem properties** | • Tainton et al. (1978) Long-term effects of burning and mowing on tall grassveld in Natal: Dry matter production.<br>• Abdalla et al. (2016) - Effects of annual burning on CO<sub>2</sub> emissions from soil.<br>• Abdalla et al. (2021) - Long-term impact of annual burning on SOC and nitrogen.<br>• Fynn et al. (2003) - Burning and its effects on soil organic matter.<br>• Melzer et al. (2010) - Fire and grazing impacts on silica production and storage.<br>• Nicolay et al. (2024) - Fire suppression and its role in maintaining soil carbon fractions.<br>• Savage and Vermeulen (1983) - Microclimate effects of spring burning.<br>• Titshall et al. (2000) - Effects of fire and herbivory exclusion on soil and vegetation.<br>• Ward et al. (2017) - Soil respiration responses to nitrogen fertilisation. | **9** |
| **Effects on plant diversity and community composition** | • Fynn et al. (2004) - Burning and mowing effects on grass and forb diversity.<br>• Fynn et al. (2005) - Long-term effects of burning and mowing on plant composition.<br>• Uys et al. (2004) - Fire regime effects on plant diversity in South African grasslands.<br>• Ward et al. (2023) - Long-term effects of burning and mowing on $\beta$ diversity. | **4** |
| **Invertebrate and microbial responses** | • Chambers and Samways (1998) - Grasshopper response to burning and mowing.<br>• Khoza et al. (2023) - Ant community responses to long-term burning and mowing.<br>• Vermeire et al. (2021) - Fire and herbivory effects on fungal and bacterial communities. | **3** |
| **Comparative studies across regions** | • Buis et al. (2009) - Controls of aboveground net primary production in mesic savanna grasslands: an inter-hemispheric comparison.<br>• Forrestel et al. (2014) - Phylogenetic and functional responses to fire in North America and South Africa.<br>• Gold et al. (2023) - Herbaceous vegetation responses to experimental fire in savannas and forests depend on biome and climate.<br>• Kirkman et al. (2014) - Divergent fire responses in South African and North American grasslands.<br>• Knapp et al. (2006) - Convergence and contingency in production–precipitation relationships in North American and South African C4 grasslands.<br>• Smith et al. (2016) - Convergence and divergence in grassland ecosystem responses to fire and grazing.<br>• Ward et al. (2020) - Are there common assembly rules for different grasslands? Comparisons of long-term data from a subtropical grassland with temperate grasslands. | **7** |
| **Remote sensing and biomass estimation** | • Sibanda et al. (2017a) - WorldView-3 capabilities in discriminating grassland management treatments.<br>• Sibanda et al. (2017b) - Estimating biomass in native grass under complex management. | **2** |

**Table 2:** Papers produced from the VFT grouped by general topic.

| TOPIC | STUDIES | NO. |
| --- | --- | --- |
| **Effects on soil and ecosystem properties** | • Buthelezi et al. (2022) - Effects of long-term (70 years) nitrogen fertilization and liming on carbon storage in water-stable aggregates of a semi-arid grassland soil.<br>• Scott and Booysen (1956) Effects of certain fertilizers on veld at Ukulinga.<br>• Ward et al. (2017b) - Soil respiration declines with increasing nitrogen fertilization and is not related to productivity in long-term grassland experiments. | **3** |
| **Effects on plant diversity and community composition** | • Barnes et al. (1987) - Fertilization of southern tall grassveld of Natal: Effects on botanical composition and utilization under grazing.<br>• Fynn and O'Connor (2005) - Determinants of community organization of a South African mesic grassland.<br>• Grunow et al. (1970) - Long term nitrogen application to veld in South Africa.<br>• Le Roux and Mentis (1986) - Veld compositional response to fertilization in the tall grassveld of Natal.<br>• Tsvuura and Kirkman (2013) - Yield and species composition of a mesic grassland savanna in South Africa are influenced by long-term nutrient addition.<br>• Zama et al. (2023) - Assessing long-term nutrient and lime enrichment effects on a subtropical South African grassland. | **6** |
| **Invertebrate and microbial responses** | • Ndabankulu et al. (2022) - Soil microbes and associated extracellular enzymes largely impact nutrient bioavailability in acidic and nutrient poor grassland ecosystem soils.<br>• Schleuss et al. (2019) - Stoichiometric controls of soil carbon and nitrogen cycling after long-term nitrogen and phosphorus addition in a mesic grassland in South Africa.<br>• Schleuss et al. (2020) - Interactions of nitrogen and phosphorus cycling promote P acquisition and explain synergistic plant-growth responses.<br>• Sithole et al. (2021a) - Nitrogen source preference and growth carbon costs of *Leucaena leucocephala* (Lam) de Wit saplings in South African grassland soils.<br>• Sithole et al. (2021b) - Altering nitrogen sources affects growth carbon costs in *Vachellia nilotica* growing in nutrient-deficient grassland soils.<br>• Tsvuura et al. (2017) - Nutrient addition increases biomass of soil fungi: evidence from a South African grassland.<br>• Zeglin et al. (2007) Microbial responses to nitrogen addition in three contrasting grassland ecosystems. | **7** |
| **Comparative studies across regions** | • Swemmer et al. (2007) - Intra-seasonal precipitation patterns and above-ground productivity in three perennial grasslands.<br>• Ward et al. (2017a) - An African grassland responds similarly to long-term fertilization to the Park Grass experiment. | **3** |
|  | • Ward et al. (2020) Are there common assembly rules for different grasslands? Comparisons of long-term data from a subtropical grassland with temperate grasslands. |  |
| **Remote sensing and biomass estimation** | • Naicker et al. (2023) - The Detection of nitrogen saturation for real-time fertilization management within a grassland ecosystem.<br>• Naicker et al. (2024) - Estimating high-density aboveground biomass within a complex tropical grassland using Worldview-3 imagery.<br>• Sibanda et al. (2015a) - Exploring the potential of in situ hyperspectral data and multivariate techniques in discriminating different fertilizer treatments in grasslands.<br>• Sibanda et al. (2015b) - Examining the potential of Sentinel-2 MSI spectral resolution in quantifying above-ground biomass across different fertilizer treatments.<br>• Sibanda et al. (2017a) - Testing the capabilities of the new WorldView-3 space-borne sensor's red-edge spectral band in discriminating and mapping complex grassland management treatments. | **5** |

Studies from the BMT and VFT both examined long-term grassland dynamics but focused on different ecological drivers. BMT research primarily investigated fire and mowing, with an emphasis on soil and ecosystem properties (nine studies) and comparative fire-driven responses across regions. In contrast, VFT studies focused on nutrient addition, exploring its effects on plant diversity (six studies) and microbial interactions (seven studies). BMT research examined the impact of fire on soil carbon, nitrogen, and microclimate, while VFT studies assessed nutrient-driven changes in soil aggregation and respiration. Both trials incorporated long-term data and remote sensing, though VFT had more studies on biomass estimation (five vs. two).

Despite their different approaches, both experiments advanced understanding of soil and ecosystem responses. Investigations on the BMT explored fire-driven changes in carbon storage, silica production, and microclimate, whereas VFT research focused on how nutrient addition altered soil respiration and microbial activity. A notable distinction lies in invertebrate and microbial studies—BMT included three studies on grasshoppers, ants, and microbial communities under fire regimes, while VFT had seven studies emphasising microbial nutrient cycling and plant-microbe interactions.

Remote sensing was applied to assess vegetation structure, composition, and biomass at both sites, with more parameters measured and technologies tested in the VFT (five studies) than the BMT (two studies).

Comparisons between these LTEs and across regions have yielded global insights. Seven BMT publications examined grassland ecosystems across different regions, focusing on productivity, fire responses, grazing effects, and community assembly rules, highlighting ecological similarities and differences between South African, North American, temperate, and subtropical grasslands. Three VFT-based papers explored grassland responses to precipitation patterns, fertilisation, and long-term ecological processes, revealing common productivity responses and assembly rules across temperate and subtropical grasslands.

### Journal and authorship distribution

Both experiments have a strong international presence. BMT research produced 22 papers in international journals and three in journals published in South Africa, while VFT research yielded 19 publications in international journals and five in locally produced journals.

Scientific findings from both experiments were published across a total of 36 venues. The most frequently used journals, appearing more than once, included the Journal of Vegetation Science (4), Soil Biology and Biochemistry (4), the African Journal of Range & Forage Science (3), Applied Vegetation Science (2), the International Journal of Remote Sensing (2), Plants (2), the Proceedings of the Annual Congresses of the Grassland Society of Southern Africa (2), and the South African Journal of Plant and Soil (2). The journals publishing the results of these two long-term experiments reflect the interdisciplinary nature of the studies, spanning vegetation science, soil ecology, remote sensing, and regional agricultural applications, with a particular focus on grassland ecosystems and their management. This broad scope indicates that the findings are relevant to various facets of ecological and environmental sciences, from specific plant and soil studies to broader topics like biodiversity, conservation, and global change biology.

In the BMT, 93 authors contributed to the papers with a diverse international mix. South Africa led with 48 main authors and co-authors (51.61%), followed by the USA with 30 authors (approximately 32.26%). Other countries contributed smaller numbers: Australia and France each had 4 authors, Brazil had 2, and Argentina, Burkina Faso, Germany, the Netherlands, and the UK each had 1 author. In contrast, the VFT experiment involved 48 authors with a much stronger local focus. South African authors dominated with 36 contributors (75%), while the international contributions were considerably lower, with Germany providing 6 authors (12.5%), the USA 5 authors (10.42%), and Australia just 1 author (2.08%). Compared to the BMT experiment, the VFT study demonstrates a more concentrated domestic involvement with only modest international participation.

The most prolific author in both the BMT and VFT experiments was KP Kirkman (from South Africa), with 11 and 10 publications, respectively (Table 3). South African researchers are the predominant authors of papers from both LTes, including notable contributions from Z Tsvuura (8 publications in the VFT) and RWS Fynn (6 in the BMT). International authors such as A Knapp and MD Smith from the USA contributed to BMT research while D Ward published papers from both experiments. Other contributors with at least two main-author publications included K Abdalla and M Sibanda for the BMT, while R Naicker, PM Schleuss and N Sithole were additional key authors for VFT.

**Table 3:** The most prolific main authors of papers from the BMT and VFT, with counts of their total and main authorship.

| **BMT** |  |  |  | **VFT** |  |  |  |
| --- | --- | --- | --- | --- | --- | --- | --- |
| **AUTHOR** | **FREQUENCY** | **MAIN AUTHOR** | **COUNTRY** | **AUTHOR** | **FREQUENCY** | **MAIN AUTHOR** | **COUNTRY** |
| Kirkman KP | 11 | 1 | SA | Kirkman KP | 10 | 0 | SA |
| Fynn RW | 6 | 3 | SA & Botswana | Tsvuura Z | 8 | 2 | SA |
| Morris CD | 6 | 0 | SA | Mutanga O | 5 | 0 | SA |
| Smith MD | 5 | 1 | USA | Magadlela A | 4 | 0 | SA |
| Knapp AK | 4 | 1 | USA | Sibanda M | 3 | 3 | SA |
| Ward D | 3 | 3 | SA & USA | Ward D | 3 | 3 | SA & USA |

### Publication citations

The distributions of citations of papers produced from the two experiments had a similar general shape (K-S test: D = 0.2283; p = 0.279), being right-skewed because of a preponderance of publications having low citation numbers and a few highly cited ones (Figure 3). Papers with 10 or fewer citations were published from 2021 onwards, except for Scott and Booysen (1956) and Tsvuura et al. (2017) from the VFT. Publications from the BMT with 100 or more citations have had time to accumulate citations since being published in 2004 or earlier (Table 4). In contrast, six of the 10 most cited VFT papers were produced in the last decade, with three of those already accumulating >100 citations (Table 5).

**Figure 3:**
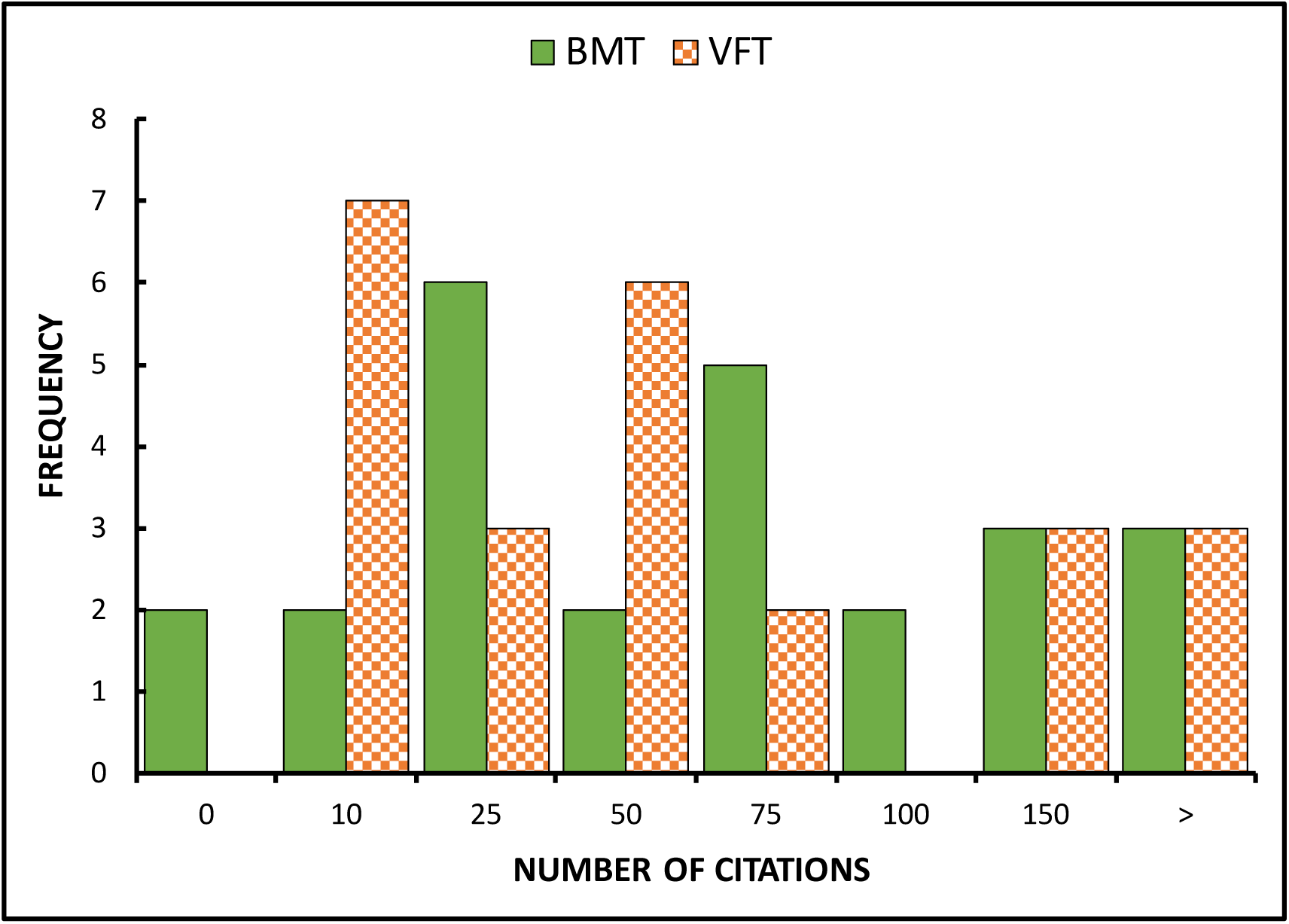
Distribution of Google Scholar citations of peer-reviewed papers produced from the Burning and Mowing Trial (BMT) and Veld Fertiliser Trial (VFT) at Ukulinga, South Africa.

**Table 4:** The top ten most cited papers from the BMT.

| NO. | AUTHORS | TITLE | CITED |
| --- | --- | --- | --- |
| 1 | Uys et al. (2004). | The effect of different fire regimes on plant diversity in southern African grasslands. | 244 |
| 2 | Fynn and Haynes (2003). | Burning causes long-term changes in soil organic matter content of a South African grassland. | 192 |
| 3 | Fynn et al. (2004). | Effect of burning and mowing on grass and forb diversity in a long-term grassland experiment. | 162 |
| 4 | Chambers and Samways (1998). | Grasshopper response to a 40-year experimental burning and mowing regime, with recommendations for invertebrate conservation management. | 130 |
| 5 | Titshall et al. (2000). | Effect of long-term exclusion of fire and herbivory on the soils and vegetation of sour grassland. | 105 |
| 6 | Knapp et al. (2006). | Convergence and contingency in production–precipitation relationships in North American and South African C4 grasslands. | 105 |
| 7 | Fynn et al. (2005). | Long-term compositional responses of a South African mesic grassland to burning and mowing. | 80 |
| 8 | Sibanda et al. (2017). | Estimating biomass of native grass grown under complex management treatments using WorldView-3 spectral derivatives. | 79 |
| 9 | Forrestel et al. (2014). | Convergent phylogenetic and functional responses to altered fire regimes in mesic savanna grasslands of North America and South Africa. | 74 |
| 10 | Melzer et al. (2010). | Fire and grazing impacts on silica production and storage in grass-dominated ecosystems. | 66 |

**Table 5:** The top ten most cited papers from the VFT.

| NO. | AUTHORS | TITLE | CITED |
| --- | --- | --- | --- |
| 1 | Swemmer et al. (2007). | Intra-seasonal precipitation patterns and above-ground productivity in three perennial grasslands. | 250 |
| 2 | Zeglin et al. (2007). | Microbial responses to nitrogen addition in three contrasting grassland ecosystems. | 239 |
| 3 | Sibanda et al. (2015b). | Examining the potential of Sentinel-2 MSI spectral resolution in quantifying above ground biomass across different fertilizer treatments. | 237 |
| 4 | Schleuss et al. (2020). | Interactions of nitrogen and phosphorus cycling promote P acquisition and explain synergistic plant-growth responses. | 106 |
| 5 | Fynn and O'Connor (2005). | Determinants of community organization of a South African mesic grassland. | 102 |
| 6 | Schleuss et a. (2019). | Stoichiometric controls of soil carbon and nitrogen cycling after long-term nitrogen and phosphorus addition in a mesic grassland in South Africa. | 102 |
| 7 | Ward et al. (2017). | Soil respiration declines with increasing nitrogen fertilization and is not related to productivity in long-term grassland experiments. | 65 |
| 8 | Sibanda et al. (2015a). | Exploring the potential of in situ hyperspectral data and multivariate techniques in discriminating different fertilizer treatments in grasslands. | 52 |
| 9 | Grunow et al. (1970). | Long term nitrogen application to veld in South Africa. | 43 |
| 10 | Sibanda et al. (2017). | Testing the capabilities of the new WorldView-3 space-borne sensor's red-edge spectral band in discriminating and mapping complex grassland management treatments. | 42 |

The papers on BMT have garnered 1,652 citations, surpassing the 1,440 citations of VFT publications. Publications from the BMT have a higher H-index (18) and i10-index (22) compared to those from the VFT, with an H-index of 16 and an i10-index of 17. Publications from the BMT have a slightly higher average citation count of 65.0, compared to 60.0 for VFT, while the median citation count is 59 for BMT and 30.5 for VFT. The higher mean and median citation counts of BMT papers reflect a larger number of mid-to-high citation papers and relatively fewer low-citation ones, while the mean for the VFT papers is biased upwards by a few highly cited works, while a preponderance of low citation publications lowers the median.

Three of the most highly referenced papers (>200 citations) from the LTEs compared multiple grassland ecosystems with varying climate and soil conditions to explore how fire, precipitation, and nitrogen influence plant diversity, productivity, and microbial activity (Tables 4 & 5). Uys et al. (2004) used data from the BMT alongside two other long-term burning experiments in semi-arid and montane grasslands in South Africa to examine forb responses to fire regimes, while Swemmer et al. (2007) investigated precipitation-productivity relationships across three sites, including the VFT, with varying annual rainfall. Zeglin et al. (2007) conducted a broader study, assessing microbial responses to increased nitrogen availability by measuring extracellular enzyme activity in soils from three grasslands in the USA (Sevilleta and Konza Prairie) and South Africa (BMT), each with contrasting edaphic and climatic conditions.

Other highly cited comparative research from the BMT (Table 4) includes studies by Titshall et al. (2000), Knapp et al. (2006) and Forrestel et al. (2014) (Table 4). Ward et al. (2007) examined soil respiration responses to two treatments in the LTEs alongside those from the Nutrient Network (NutNet) on the Ukulinga plateau. In an early cross-site assessment, Grunow et al. (1970) evaluated the effects of fertiliser addition across multiple grasslands, including the VFT, in South Africa. These multi-site studies have had broad appeal by providing key insights into ecological processes across diverse ecosystems, advancing our understanding of how environmental factors shape biodiversity and ecosystem function globally.

Remote sensing studies on both LTEs (Tables 4 & 5) have also been especially popular for their insights into cost-effective satellite sensing technology and data processing methods that can enable rapid, large-scale assessments to detect fine-scale vegetation variation in other grassland ecosystems.

### Wider citation patterns

Quantitative citation data from Scopus reveal distinct patterns of academic engagement and scholarly influence between the two experiments. The BMT publications accumulated 806 scholarly citations from 560 different principal authors, while papers from the VFT experiment generated 931 citations from 692 authors. The citation distributions exhibited a heavy-tailed pattern, wherein a small cohort of researchers contributed disproportionately to the total citations, while a much larger number of authors cited works only once or twice. Single citations represented 78.0% and 81.9% of all citations for the BMT and VFT, respectively. The top five authors that most frequently cited BMT publications were CD Morris (13), D Ward (10), RWS Fynn (10), L Joubert (8), and SE Koerner (8) while and those authors who referred to VFT papers most often were M Sibanda (19), T Dube (11), C Shoko (11), R Naicker (8), and J Li (7).

The source publications citing the LTE research were also similarly skewed. Citations of the most prominent BMT research were distributed across 290 journals, with a concentrated presence in a few specialised rangeland and ecological publications such as the African Journal of Range and Forage Science (72), Applied Vegetation Science (23), Journal of Vegetation Science (21), Journal of Ecology (19), Biodiversity and Conservation (16), and South African Journal of Botany (15), cumulatively contributing more than 20% of all the citations to BMT work. Journals citing the top 10 VFT papers spanned a more interdisciplinary set of 282 journals focused on technological applications (such as remote sensing), soil science, and environmental change. There was a prominent representation of citations (>20%) in Remote Sensing (56), Soil Biology and Biochemistry (39), Global Change Biology (27), Plant and Soil (27), African Journal of Range and Forage Science (20), and Science of the Total Environment (19).

Citations of the BMT’s top papers originated from principal authors based in 59 countries, with South Africa (231) accounting for more than 28%, followed by the USA (187), China (75), Australia (42), Germany (39), and the UK (25), collectively making up almost 75% of all citations. The most-referenced VFT papers were cited by main authors from 63 countries, with the highest counts from China (383; 41.2%), South Africa (153), the USA (121), and Germany (43), together accounting for three-quarters of the total citations.

Overall, while both experiments command substantial global attention, BMT’s impact is anchored in niche rangeland and grassland ecological fields, whereas VFT demonstrates broader appeal through its integration into diverse technological and environmental research communities.

## DISCUSSION

Over the past 75 years, the very long-term burning, mowing and nutrient addition experiments on grassland at Ukulinga, Pietermaritzburg, have yielded a modest scientific output, with just over three peer-reviewed papers per decade. However, output accelerated since 2000, with 87.5% of BMT publications and 83.3% of VFT publications produced in the past quarter century. Moreover, research from the Ukulinga LTEs has had broad international reach, with the top 10 cited papers from each experiment collectively cited in 458 journals by 1172 principal authors from 78 countries outside South Africa. The growing interest in these long-term field experiments, both locally and internationally, illustrates the importance of not only conducting research over extended periods with consistent treatment application but also ensuring their long-term maintenance. Such enduring outdoor laboratories enable researchers to not only track temporal trends but to pursue new, often multidisciplinary, scientific investigations—many of which were not originally envisaged—and to explore interactions with emerging environmental pressures, such as those driven by anthropogenic climate change (Leigh and Johnston 1994).

Several comparative studies between the two experiments at Ukulinga and with those elsewhere in South Africa (Kruger National Park), the USA, and the UK have been published (see Tables 1 & 2), testing general hypotheses and providing useful ecological insights. However, other defunct and ongoing long-term experiments in southern Africa could also provide valuable contrasts to those at Ukulinga. Two notable extant long-term burning experiments are: (1) the Experimental Burn Plot (EBP) trial, established in 1954 in Kruger National Park’s subtropical savannas to study fire regimes across landscapes and inform fire management (Biggs et al. 2003; van Wilgen et al. 2007); and (2) the Brotherton Burning Trial (BBT), initiated in 1980 at Cathedral Peak to test burn frequency and seasonality in replicated plots (Short 2007; Morris et al. 2021). Several discontinued long-term burning experiments have also been conducted elsewhere in Africa (Swift et al. 1994), including in the Matopos savanna in Zimbabwe (O’Connor 1985; Furley et al. 2008) and in semi-arid grassland at Fort Hare, Alice, South Africa (Oluwole et al. 2008).

There are also opportunities to compare nutrient addition effects in the VFT with other fertiliser trials, apart from the Park Grass Experiment—the oldest of all—established in 1856 on species-rich pasture at Rothamsted, England (Silvertown et al. 2006; Ward et al. 2017a). Several grassland fertiliser trials were launched in South Africa in the 1940s, aiming to repurpose surplus explosives chemicals and improve natural grassland by adding mineral nutrients. Most were soon abandoned when it became clear that fertilisation, while boosting productivity, caused undesirable shifts in species composition and reduced forage quality (Roux and Roux 1969; Grunow 1970). One exception was a savanna trial at Towoomba in north-eastern South Africa, which ran from 1948 until 1995 (Mills et al. 2017).

Below, we discuss how the Ukulinga LTEs have: (1) served as valuable field experiments generating novel insights for grassland ecology and rangeland management; (2) played a key role in international collaborative research across countries and continents; and (3) supported formal and informal education since their inception. We also reflect on lessons learned from managing long-term field experiments and explore ways to enhance their value in the future.

### The Ukulinga LTEs as an outdoor research laboratory

The diverse scientific outputs from the Ukulinga long-term experiments, spanning molecular, microscopic, macroscopic, and community-level scales, and covering both biotic and abiotic components above and below ground, have steadily contributed to the growing body of basic knowledge on grassland ecosystems and their management. Many findings have reinforced results from other regions, whereas some have been foundational, revealing novel insights that have shaped current understanding and management of mesic grasslands in southern Africa, particularly regarding the role of disturbance by fire.

The BMT and VFT differ fundamentally in two key ways: the type of disturbance applied and how they exert their effect on the grassland ecosystem. First, fire and defoliation can be considered natural disturbances, integral to the evolution and resilience of grasses (Strömberg 2011; Linder et al. 2018) and the maintenance of open grasslands (Bond 2019), whereas nutrient addition through repeated applications of concentrated mineral fertiliser has no analogue in uncultivated grasslands, not even in rangelands exposed to atmospheric nitrogen and other pollutants (Stevens et al. 2022). Second, in the BMT, the removal of periodic disturbance by fire and mowing had the greatest influence on the grassland’s structure, composition, and state. In contrast, in the VFT, the regular addition of different types and levels of mineral nutrients primarily drove changes in the grassland.

Decades of fire and mowing exclusion transformed the original open grassland at Ukulinga into a dense, woody-dominated system. Woody plant density exceeded 10,000 individuals per hectare (some with a > 0.5 m girth), with alien invasive species such as *Acacia mearnsii, Jacaranda mimosifolia*, *Melia azedarach,* and the shrub, *Lantana camara,* forming much of the cover, alongside some indigenous microphyllous and broadleaf trees (Titshall et al. 2000). The grass layer was dominated by shade-tolerant species, while grasses abundant in the defoliated plots were absent (Fynn et al. 2005). Grass species richness declined markedly, likely due to litter accumulation and shading, whereas forb richness was maintained but with a major shift toward shade-tolerant species (Fynn et al. 2004). This restructured plant community reflects a state shift from an open, disturbance-maintained grassland to a self-reinforcing woody system driven by the long-term absence of fire and defoliation (Wilcox et al. 2018; Pausas and Bond 2020). These changes mirror patterns seen in other protected mesic grasslands in South Africa (Westfall et al. 1983; Titshall et al. 2000) and globally under suppressed disturbance regimes (Ratajczak et al. 2014; Bond 2019).

Such a major shift in vegetation state did not occur with long-term nutrient addition in the VFT, which remained a grassland without the ingress of woody species over 75 years (Zama et al. 2023). Instead, fertilisers altered sward structure, productivity and species composition, and reduced plant species diversity, a pattern consistent with findings from grasslands elsewhere (Borer et al. 2017; You et al. 2017; Shi et al. 2024). Nutrient addition, especially high levels of N, increased productivity and sward height while driving compositional shifts (le Roux and Mentis 1986; Barnes et al. 1987; Tsvuura and Kirkman 2013). Taller, fast-growing, nitrophilous grasses were favoured by additional nutrients over the original shorter dominants (Fynn and O’Connor 2005), mirroring the effects of heavy grazing (le Roux and Mentis 1986; Barnes et al. 1987). Lime had little impact on productivity but influenced species composition through changes in soil pH, which was strongly determined by the type of N-fertiliser applied (Zama et al. 2023). The effects of long-term nutrient addition on soil chemical and physical properties in the VFT remain to be fully described, but studies suggest that shifts in microbial communities and their activity may be key mechanisms driving changes in soil nutrient dynamics and availability to plants (e.g., Zeglin et al. 2007; Schleus et al. 2019, 2020; Ndabankulu et al. 2022).

Apart from revealing the main dynamics in response to disturbance, briefly summarised above, the two LTEs at Ukulinga have been particularly useful owing to their replicated factorial design, allowing detection of important interactions between the type, timing and intensity of disturbance on various grassland characteristics. In the BMT, for example, species composition in plots mown during the growing season was little affected by when and how often they were burned during the dormant period. However, in plots not mown in summer, burning frequency and season did alter species abundance, favouring taller grass dominants in triennially burned plots (Fynn et al. 2005). The sequence of defoliation also proved important, with species richness most affected by the interaction between summer mowing and dormant-season burning. For example, winter burning had little impact following early summer mowing but led to a marked decline in richness after late summer mowing, possibly due to greater soil desiccation (Fynn et al. 2004). The most striking interaction in the VFT was that *Eragrostis curvula* came to dominate the high-N plots, but when P was also added, *Panicum maximum* prevailed (O’Connor and Fynn 2005). There were also complex three-way interactions between LAN, phosphorus, and lime affecting forb richness that warrant further study, with lime increasing forb richness at high LAN levels only when phosphorus was absent (Zama et al. 2023). Such contingencies combined with strong main disturbance effects have created a mosaic of grassland communities and a patchwork of varied sward structures and compositions that provide a valuable setting to study fauna, microbes, and remote sensing methods (Tables 1&2).

### LTEs for comparative studies and networking

Further value has been extracted from the two Ukulinga LTEs by contrasting their disturbance responses with other experiments, locally and abroad. For example, Ward et al. (2017a, 2017b, 2020) showed that comparing local (Ukulinga) and international long-term experiments reveals consistent ecosystem responses, such as nitrogen reducing soil respiration and species richness, and that key ecological patterns, including burning, fertilisation, and soil chemistry effects, emerge only over long timescales and are governed by shared community assembly rules across climates. Cross-continental contrasts with the BMT found global productivity drivers consistent but forb responses regionally variable (Buis et al. 2009); fire caused convergent phylogenetic and functional grassland responses through consistent ecological filtering across distinct biogeographic histories (Forrestel et al. 2014); shared mechanisms led to convergent ecosystem functioning under fire and grazing despite divergent plant-community responses (Smith et al. 2016); and similar climates produced comparable productivity–precipitation patterns but revealed region-specific seasonal sensitivities (Knapp et al. 2006). Using VFT data, Swemmer et al. (2007) showed that precipitation amount, event size, frequency, and timing affected productivity differently across three southern African grasslands, with intra-seasonal rainfall patterns influencing productivity beyond total rainfall.

Although local researchers, most notably KP Kirkman from UKZN, have been the most prolific authors of publications from the Ukulinga LTEs, many of the comparative studies mentioned above have involved international teams, often led by US principal investigators. Collaboration between local and US researchers began in the mid-2000s, with early examples including Knapp and coauthors’ study on production and precipitation relationships across the USA and South Africa (Knapp et al. 2006) and Zeglin et al.’s (2007) research on microbial responses to nitrogen addition in the VFT. The isolation of local researchers during the Apartheid era initially limited their connections, but two articles in non-technical publications (Morris 2001, 2002) helped raise international awareness, particularly in the USA, of the Ukulinga LTEs and their research potential, fostering the collaboration that has emerged this century.

Informal networking is likely to remain the primary avenue for future collaborations involving the Ukulinga LTEs as there is no dedicated funding beyond basic operational costs and the site is not part of a formal framework such as the Long-Term Ecological Research network of sites (Knapp et al. 2012). However, the establishment of nearby medium-term experiments, including NutNet (since 2009; www.nutnet.org), DroughtNet (since 2019; www.droughtnet.weebly.com) and DragNet (since 2020; www.dragnetglobal.org), adjacent to the VFT, may support efforts to formalise the Ukulinga plateau as a long-term grassland research site for both research and education.

### LTEs for demonstration and education

The impact of the Ukulinga LTEs through education is not easily quantified but several of the papers published on the experiments were based on MSc and PhD studies done at UKZN. Examples include publications from the BMT, such as Chambers and Samways (1998), Uys et al. (2004), Fynn et al. (2003), and Nicolay et al. (2024), as well as Zama et al. (2023) from the VFT. Some other unpublished postgraduate theses undertaken on the two trials were Booysen (1954), Pienaar (1956), I’Ons (1960), Edwards (1961), Dillon (1980), and Demmer (2019). Over the years, many UKZN undergraduates have done field practicals at Ukulinga, sometimes resulting not just in sore knees and aching backs (Morris and Fynn 2001), but in BSc Honours or 4th-year project reports, such as Rodel (1950), Dickens (1951), Routledge (1951), Thompson (1994), Webster (1996a), Webster (1996b), and Van Wyk (1998).

Overall, the ‘Natal School’ of grassland scientists and ecologists at the University of Natal/KwaZulu-Natal, which initiated and runs the Ukulinga LTEs, has had a prominent influence on how fire is viewed and advocated as a key tool for sustainable grassland management in South Africa (Pooley 1998, 2022). This dogma has been strongly advocated through textbooks (Tainton 1981, 1999) and other management guides (Danckwerts and Teague 1989; Engelbrecht et al. 2004; SANBI 2014; Van Oudtshoorn 2015). Many undergraduate and postgraduate students trained at the university, exposed to the LTEs and their findings, emerged as critics of grassland fertilisation and as committed ‘pyrophiles’ who believe that mesic grasslands (sourveld) “owe their existence to fire” (Tainton 1981, 1999). The most renowned of these was Winston Trollope, who went on to have a major influence across the subcontinent in promoting fire as an indispensable management tool for grasslands and savannas (Govender et al. 2022).

The Ukulinga LTEs have also welcomed numerous visitors, including students, academics, extension officers, conference attendees, and both formal and informal conservation organisations. These groups have observed and discussed the notable treatment effects on grassland ecology and management, and may have helped to inform policy, advice, and legislation on sustainable grassland management.

### Que Vadis, the Ukuling LTEs?

The original treatments in the BMT and VFT have been maintained for nearly 75 years, with only minor adjustments. In the BMT, the summer mowing that was initially combined with autumn burns was dropped early, owing to insufficient fuel to implement an autumn burn. A seven-yearly no-mow test year to measure the effect of all treatments (except the protected control) on herbage production was later suspended after 1997 to allow more realistic vegetation development under infrequent burning; since then, saplings and aloes have begun to invade (Nkabinde 2020). The seven-yearly harvests and annual mowing offtakes showed that spring burns applied after regrowth had begun temporarily reduced yield compared to winter mowing (Tainton et al. 1978), but biennial burning enhanced average herbage production and forage quality by promoting nutritious regrowth and maintaining palatable grasses (Tainton et al. 1978; Mentis and Tainton 1984; Tainton and Mentis 1984). One fire exclusion plot was accidentally burned in 1958.

A deliberate modification in the VFT was the decision in 2019 to stop fertiliser application on a random half of each plot, while maintaining the original treatment on the other half. This was done to assess whether changes in vegetation and soils, particularly acidification from long-term nutrient addition, could be reversed, and to monitor the rate and direction of recovery. A similar division to study reversals under more frequent burning in infrequently burned plots would not be possible in the BMT, given the already limited plot size available for fire research (Swift et al. 1994). Minor adjustments to improve implementation, as noted above, are common in LTEs. However, changes to the original treatments should be avoided, even if initial objectives appear met and research questions answered. This is because some effects take time to emerge, unexpected changes may occur, and new questions can arise that are best addressed using data from or further multidisciplinary and interdisciplinary studies on the original experiments (Leigh and Johnson 2004). Furthermore, progressive vegetation succession, seasonal cycles, tipping points, and state changes can only be revealed through long-term study (Lindenmayer et al. 2012; Peterson et al. 2012). Perhaps most valuable is the understanding gained from deeply studying a single ecosystem for a long time (Riginos et al. 2024). We therefore strongly recommend that the long-term burning and mowing, and nutrient addition experiments at Ukulinga be maintained without alteration for as long as possible.

## CONCLUSION

The very long-term experiments at Ukulinga, which have run uninterrupted for 75 years, represent a rare and valuable resource for understanding grassland dynamics under sustained disturbance regimes in southern Africa. Their longevity has enabled the detection of slow ecological responses and important treatment interactions often overlooked in shorter studies, producing insights that have informed sustainable grassland management. Over time, like the long-running Grass Park Study at Rothamsted, the Ukulinga experiments have become, in the words of J.B. Lawes (Johnston 1994), a “battlefield of the plants” (and of the micro- and meso-fauna), all vying for resources and space under different disturbance regimes. This dynamic has drawn the interest of a wide array of plant, soil, zoological, and other scientists, well beyond the field of agriculture. The multi-disciplinary studies conducted on the BMT and VFT have contributed to ecological theory, shaped local management practices, and supported a wide range of comparative research, networking, and education. In the context of ongoing climate and environmental change, maintaining these experiments provides a stable reference point for monitoring ecological responses, testing new hypotheses, and supporting collaborative research to address outstanding scientific questions. Ensuring their continuity will preserve the ability to detect long-term patterns and processes that only emerge over extended timescales.

