## Supplementary Table 1 for "Seventy-five years of insight – the impact of the Ukulinga very long-term grassland experiments"

**Supplementary Table S2: Peer-reviewed scientific papers from the VFT.**

| NO. | AUTHORS | YEAR | TITLE | JOURNAL | ISSUE | GOOGLE SCHOLAR CITATI |
|---|---|---|---|---|---|---|
| 1 | Barnes GR, Bransby DI, Tainton NM | 1987 | Fertilization of southern tall grassveld of Natal: Effects on botanical composition and utilization under grazing | Journal of the Grassland Society of Southern Africa | 4: 63-67. | 16 |
| 2 | Buthelezi K, Buthelezi-Dube N | 2022 | Effects of long-term (70 years) nitrogen fertilization and liming on carbon storage in water-stable aggregates of a semi-arid grassland so | Heliyon | 8(1) e08690 | 11 |
| 3 | Fynn RW, O'connor TG | 2005 | Determinants of community organization of a South African mesic grassland | Journal of Vegetation Science | 16: 93-102. | 102 |
| 4 | Grunow JO, Pienaar AJ, Breytenbach C | 1970 | Long-term nitrogen application to veld in South Africa | Proceedings of the Annual Congresses of the Grassland Society of Southern Afric | 5: 75-90. | 43 |
| 5 | Le Roux NP, Mentis M | 1986 | Veld compositional response to fertilization in the tall grassveld of Nata | South African Journal of Plant and Soi | 3: 1-10. | 26 |
| 6 | Naicker R, Mutanga O, Peerbhay K, Agjee N | 2023 | The detection of nitrogen saturation for real-time fertilization management within a grassland ecosystem | Applied Sciences | 13: 4252. | 5 |
| 7 | Naicker R, Mutanga O, Peerbhay K, Odebiri O | 2024 | Estimating high-density aboveground biomass within a complex tropical grassland using Worldview-3 imagery | Environmental Monitoring and Assessment | 196: 370. | 4 |
| 8 | Ndabankulu K, Egbewale SO, Tsvuura Z, Magadlela A | 2022 | Soil microbes and associated extracellular enzymes largely impact nutrient bioavailability in acidic and nutrient-poor grassland ecosystem soil | Scientific Reports | 12: 12601. | 32 |
| 9 | Schleuss PM, Widdig M, Heintz-Buschart A, Guhr A, Martin S, Kirkman K, Spohn M | 2019 | Stoichiometric controls of soil carbon and nitrogen cycling after long-term nitrogen and phosphorus addition in a mesic grassland in South Afric | Soil Biology and Biochemistry | 135: 294-303. | 102 |
| 10 | Schleuss PM, Widdig M, Heintz-Buschart A, Kirkman K, Spohn M | 2020 | Interactions of nitrogen and phosphorus cycling promote P acquisition and explain synergistic plant-growth response | Ecology | 101: e03003. | 106 |
| 11 | Scott JD, Booysen PdeV | 1956 | Effects of certain fertilizers on veld at Ukulinga | South African Journal of Science | 52: 240-243. | 6 |
| 12 | Sibanda M, Mutanga O, Rouget M, Odindi J | 2015 | Exploring the potential of in situ hyperspectral data and multivariate techniques in discriminating different fertilizer treatments in grassland | Journal of Applied Remote Sensing | 9: 096033-096033. | 52 |
| 13 | Sibanda M, Mutanga O, Rouget M | 2015 | Examining the potential of Sentinel-2 MSI spectral resolution in quantifying above-ground biomass across different fertilizer treatment | ISPRS Journal of Photogrammetry and Remote Sensing | 110: 55-65. | 237 |
| 14 | Sibanda M, Mutanga O, Rouget M | 2017 | Testing the capabilities of the new WorldView-3 space-borne sensor's red-edge spectral band in discriminating and mapping complex grassland management treatmen | International Journal of Remote Sensing | 38: 1-22. | 42 |
| 15 | Sithole N, Tsvuura Z, Kirkman K, Magadlela A | 2021 | Nitrogen source preference and growth carbon costs of Leucaena leucocephala (Lam) de Wit saplings in South African grassland soil | Plants | 10: 2242. | 7 |
| 16 | Sithole N, Tsvuura Z, Kirkman K, Magadlela A | 2021 | Altering nitrogen sources affects growth carbon costs in Vachellia nilotica growing in nutrient-deficient grassland soi | Plants | 10: 1762. | 3 |
| 17 | Swemmer AM, Knapp AK, Snyman HA | 2007 | Intra-seasonal precipitation patterns and above-ground productivity in three perennial grassland | Journal of Ecology | 95: 780-788. | 250 |
| 18 | Tsvuura Z, Kirkman KP | 2013 | Yield and species composition of a mesic grassland savanna in South Africa are influenced by long-term nutrient additio | Austral Ecology | 38: 959-870. | 29 |
| 19 | Tsvuura Z, Kirkman KP, Avolio ML | 2017 | Nutrient addition increases biomass of soil fungi: evidence from a South African grassland | South African Journal of Plant and Soi | 34: 71-73. | 4 |
| 20 | Ward D, Kirkman K, Tsvuura Z | 2017 | An African grassland responds similarly to long-term fertilization to the Park Grass experiment | PLoS One | 12: e0177208. | 36 |
| 21 | Ward D, Kirkman K, Hagenah N, Tsvuura Z | 2017 | Soil respiration declines with increasing nitrogen fertilization and is not related to productivity in long-term grassland experiment | Soil Biology and Biochemistry | 115: 415-422. | 65 |
| 22 | Ward D, Kirkman KP, Tsvuura Z, Morris C, Fynn RW | 2020 | Are there common assembly rules for different grasslands? Comparisons of long-term data from a subtropical grassland with temperate grassland | Journal of Vegetation Science | 31 :780-791. | 17 |
| 23 | Zama N, Magadlela A, Mkhize N, Tedder M, Kirkman K, Twine W | 2023 | Assessing long-term nutrient and lime enrichment effects on a subtropical South African grassland | African Journal of Range & Forage Science | 40: 206-218. | 6 |
| 24 | Zeglin LH, Stursova M, Sinsabaugh RL, Collins SL | 2007 | Microbial responses to nitrogen addition in three contrasting grassland ecosystems | Oecologia | 154: 349-359. | 239 |
