## Supplementary Table 2 for "Seventy-five years of insight – the impact of the Ukulinga very long-term grassland experiments"

**Supplementary Table S1: Peer-reviewed scientific papers from the BMT.**

| NO. | AUTHORS | YEAR | TITLE | JOURNAL | ISSUE | GOOGLE SCHOLAR CITATIONS |
|---|---|---|---|---|---|---|
| 1 | Abdalla K, Chivenge P, Ciais P, Chaplot V | 2021 | Long-term (64 years) annual burning lessened soil organic carbon and nitrogen content in a humid subtropical grassland. | Global Change Biology | 27: 6436-6453. | 10 |
| 2 | Abdalla K, Chivenge P, Everson C, Mathieu O, Thevenot M, Chaplot V | 2016 | Long-term annual burning of grassland increases CO2 emissions from soils. | Geoderma | 282: 80-86. | 30 |
| 3 | Buis GM, Blair JM, Burkepile DE, Burns CE, Chamberlain AJ, Chapman PL, Collins SL, Fynn RW, Govender N, Kirkman KP, Smith MD | 2009 | Controls of aboveground net primary production in mesic savanna grasslands: an inter-hemispheric comparison. | Ecosystems | 12: 982-995. | 59 |
| 4 | Chambers BQ, Samways MJ | 1998 | Grasshopper response to a 40-year experimental burning and mowing regime, with recommendations for invertebrate conservation management. | Biodiversity & Conservation | 7: 985-1012. | 130 |
| 5 | Forrestel EJ, Donoghue MJ, Smith MD | 2014 | Convergent phylogenetic and functional responses to altered fire regimes in mesic savanna grasslands of North America and South Africa. | New Phytologist | 203: 1000-1011. | 74 |
| 6 | Fynn RW, Haynes RJ, O'connor TG | 2003 | Burning causes long-term changes in soil organic matter content of a South African grassland. | Soil Biology and Biochemistry | 35: 677-687. | 192 |
| 7 | Fynn RW, Morris CD, Edwards TJ | 2004 | Effect of burning and mowing on grass and forb diversity in a long-term grassland experiment. | Applied Vegetation Science | 7: 1-10. | 162 |
| 8 | Fynn RW, Morris CD, Edwards TJ | 2005 | Long-term compositional responses of a South African mesic grassland to burning and mowing. | Applied Vegetation Science | 8: 5-12. | 80 |
| 9 | Gold ZJ, Pellegrini AFA, Refsland TK, Andrioli RJ, Bowles ML, Brockway DG, Burrows N, Franco AC, Hallgren SW, Hobbie SE, Hoffmann WA, [illegible] | 2023 | Herbaceous vegetation responses to experimental fire in savannas and forests depend on biome and climate. | Ecology Letters | 26: 1237-1246. | 15 |
| 10 | Khoza LR, Andersen AN, Munyai TC | 2023 | Effect of long-term burning and mowing regimes on ant communities in a mesic grassland. | Diversity | 15(9): 996. | 0 |
| 11 | Kirkman KP, Collins SL, Smith MD, Knapp AK, Burkepile DE, Burns CE, Fynn RW, Hagenah N, Koerner SE, Matchett KJ, Thompson DI | 2014 | Responses to fire differ between South African and North American grassland communities. | Journal of Vegetation Science | 25: 793-804. | 61 |
| 12 | Knapp AK, Burns CE, Fynn RW, Kirkman KP, Morris CD, Smith MD. | 2006 | Convergence and contingency in production–precipitation relationships in North American and South African C4 grasslands. | Oecologia | 149: 456-64. | 105 |
| 13 | Melzer SE, Knapp AK, Kirkman KP, Smith MD, Blair JM, Kelly EF | 2010 | Fire and grazing impacts on silica production and storage in grass-dominated ecosystems. | Biogeochemistry | 97: 263-278. | 66 |
| 14 | Nicolay RE, Mkhize NR, Tedder MJ, Kirkman KP | 2024 | Fire suppression interacts with soil acidity to maintain stable recalcitrant pyrogenic carbon fractions in South African mesic grassland soils. | African Journal of Range & Forage Science | 21: 1-9. | 0 |
| 15 | Savage MJ, Vermeulen K | 1983 | Microclimate modification of tall moist grasslands of Natal by spring burning. | Rangeland Ecology & Management/Journal of Range Management Archives | 36: 172-174. | 25 |
| 16 | Sibanda M, Mutanga O, Rouget M | 2017 | Testing the capabilities of the new WorldView-3 space-borne sensor’s red-edge spectral band in discriminating and mapping complex grassland management treatments. | International Journal of Remote Sensing | 38: 1-22. | 42 |
| 17 | Sibanda M, Mutanga O, Rouget M, Kumar L | 2017 | Estimating biomass of native grass grown under complex management treatments using WorldView-3 spectral derivatives. | Remote Sensing | 9(1): 55. | 79 |
| 18 | Smith MD, Knapp AK, Collins SL, Burkepile DE, Kirkman KP, Koerner SE, Thompson DI, Blair JM, Burns CE, Eby S, Forrestel EJ | 2016 | Shared drivers but divergent ecological responses: Insights from long-term experiments in mesic savanna grasslands. | BioScience | 66: 666-682. | 24 |
| 19 | Tainton NM, Booysen PD, Bransby DI, Nash RC | 1978 | Long-term effects of burning and mowing on tall grassveld in Natal: Dry matter production. | Proceedings of the Annual Congresses of the Grassland Society of Southern Africa | 13: 41-44. | 24 |
| 20 | Titshall LW, O'Connor TG, Morris CD | 2000 | Effect of long-term exclusion of fire and herbivory on the soils and vegetation of sour grassland. | African Journal of Range & Forage Science | 17: 70-80. | 105 |
| 21 | Uys RG, Bond WJ, Everson TM | 2004 | The effect of different fire regimes on plant diversity in southern African grasslands. | Biological Conservation | 118: 489-499. | 244 |
| 22 | Vermeire ML, Thoresen J, Lennard K, Vikram S, Kirkman K, Swemmer AM, Te Beest M, Siebert F, Gordijn P, Venter Z, Brunel C | 2021 | Fire and herbivory drive fungal and bacterial communities through distinct above- and below-ground mechanisms. | Science of the Total Environment | 785: 147189. | 14 |
| 23 | Ward D, Kirkman K, Hagenah N, Tsvuura Z | 2017 | Soil respiration declines with increasing nitrogen fertilization and is not related to productivity in long-term grassland experiments. | Soil Biology and Biochemistry | 115 :415-422. | 65 |
| 24 | Ward D, Kirkman K, Morris C | 2023 | Long-term subtropical grassland plots take a long time to change: Replacement is more important than richness differences for beta diversity. | Ecology and Evolution | 13: 1-17. | 2 |
| 25 | Ward D, Kirkman KP, Tsvuura Z, Morris C, Fynn RW | 2020 | Are there common assembly rules for different grasslands? Comparisons of long-term data from a subtropical grassland with temperate grasslands. | Journal of Vegetation Science | 31: 780-791. | 17 |
